# hAM-pe soluble extract increases human keratinocyte HaCaT cell-line mobility in scratch assays

**DOI:** 10.64898/2026.11.04.764050

**Authors:** Carla M. Soprano, María Ximena Guerbi, Flavia M. Michelini, Alejandro Berra, Griselda N. Moreno, Tomás S. Grigera

**Affiliations:** Instituto de Física de Líquidos y Sistemas Biológicos (IFLySiB), CONICET — Universidad Nacional de La Plata, La Plata, Argentina; CCT CONICET La Plata, Consejo Nacional de Investigaciones Científicas y Técnicas, Argentina; Centro de Medicina Traslacional, Hospital El Cruce “Dr. Néstor Carlos Kirchner” (CEMET–HEC), Florencio Varela, Argentina; Instituto de Ciencias de la Salud, Universidad Nacional Arturo Jauretche, Florencio Varela, Argentina; Instituto de Estudios Inmunológicos y Fisiopatológicos (IIFP), CONICET, La Plata, Argentina; Departamento de Física, Facultad de Ciencias Exactas, Universidad Nacional de La Plata, Argentina; Istituto Sistemi Complessi, Consiglio Nazionale delle Ricerche, UOS Sapienza, Rome, Italy

**Keywords:** human amniotic membrane, wound healing, keratinocyte migration, HaCaT cells, scratch assay, cell motility

## Abstract

We have studied the impact of human amniotic membrane (hAM-pe) on wound closing speed in a scratch assay of human keratinocytes (HaCaT). Though not without limitations, scratch assays are useful for studying collective cellular displacement as a model of wound healing. We show that the treatment is effective in decreasing the average time that the cells need to completely close the wound, in agreement with previous *in vivo* experiments. We developed an automated segmentation software, and verified that a linear fit, though not completely describing the wound edge dynamics, is a robust method for comparing treatments and control measurements. By tracking the individual motion of a subset of cells, we found that individual treatment does not cause an increase in individual speed enough to explain the effect of the treatment at the collective (wound edge) level. Instead, the mean squared displacement and numerical simulations of a simple model of interacting active Brownian particles support the view that the main effect of the treatment at the individual level is to increase the persistence of cellular motion.

**Subject Areas:** biophysics, computational biology, biomaterials

## 1. Introduction

Cell migration is an essential component of tissue repair, and depends on the coordination of biochemical signals, cell-cell interactions, and extracellular matrix properties. Cell migration studies are thus especially relevant to evaluate bio-materials for wound treatment, because they allow to explore whether those materials create conditions compatible with the displacement of the cells involved in tissue repair. Although migration is only one dimension of cicatrisation (by itself, it cannot account for the integration of inflammatory, vascular, and repair processes involved in actual wound healing), *in vitro* biological models allow to study cellular response under controlled conditions, and to compare treatments before moving on to systems of higher complexity. Keeping in mind that the choice of biological model and quantification methods constrain the information that can be obtained as well as the relevance of the conclusions reached [1, 2], the scratch assay [3] is an important tool in cell migration studies. Consisting in the examination of the closure of a cell-free region created by scratching a cell monolayer, it is a useful and accessible approximation to study collective cellular displacement. A limitation of scratch assays is that scratching to produce the cell-free region can damage cells and alter the extracellular matrix; however, other methods such as barrier exclusion avoid possible mechanical injury but do not reproduce the damage signals. Three-dimensional systems and microfluidic platforms may allow better reproduction of other characteristics of the tissue environment, but also need adequate validation [2]. Thus, notwithstanding their limitations, scratch assays are a useful probe into cell motility under injury, and complementary to other methods [1].

Furthermore, the experimental information obtained from scratch assays combined with mathematical modelling can help formulate hypotheses on the mechanisms of collective motion. Stochastic models introduce rules of local motion and cell interaction, and allow to study their consequences at the population level [4, 5].

Here we study the effect of human amniotic membrane on wound healing in a scratch assay of a spontaneously immortalized human keratinocyte line (HaCaT). Human amniotic membrane has been widely used as a biological dressing. Previously, some of us found that the healing of complex skin wounds has been surprisingly improved in patients treated with a novel homogenized, lyophilized and sterilized amniotic membrane patch (hAM-pe) [6, 7]. To elucidate the mechanisms underlying the wound-healing promotion induced by hAM-pe, we evaluate the effect of soluble extracts from homogenized and lyophilized human amniotic membrane on cell migration. We focus on wound edge speed rather than relative area, since the latter is sensitive to wound geometry [8, 9]. In addition to the closure assay, we track individual motion of a subset of dyed cells, using this information together with an agent-based mathematical model (active Brownian particles [10, 11] with excluded volume) to look for quantitative and qualitative changes in the characteristics of cellular motion under treatment.

## 2. Methods

### (a) Culture and scratch assay

Briefly, HaCaT cells were cultured in T75 flasks in optimized basal medium and detached using TrypLE™. To facilitate single-cell tracking, one-tenth of the cell suspension was fluorescently labeled with CellTrace™ Violet according to the manufacturer’s instructions and recombined with the unlabelled fraction (1:10 ratio). Cells were then seeded into 24-well plates at a density adjusted to reach near-confluent monolayers within 24 h and allowed to adhere overnight (ON) (37 ^*°*^C, 5% CO_2_) prior to the assay. To achieve cell cycle synchronization, the complete culture medium was replaced with serum-free medium ON. Serum-deprivation synchronizes cells through proliferation arrest, which, given a doubling time of 28 h [12–14], ensures that wound closure differences observed during at least the first 18 h are motogenic rather than mitogenic. Following synchronization, scratch wounds were performed, and wells were washed three times with sterile PBS to remove non-adherent cells. Culture medium supplemented with 2% FBS was then added alone or containing hAM-pe soluble extracts at the tested concentrations. The plate was transferred to a Cytation C10 incubator (37 ^*°*^C, 5% CO_2_) to monitor cell migration over 24 h. Five to six regions of interest (ROIs) were defined per well. Three independent experiments were performed. A detailed protocol will be published elsewhere [15].

### (b) Treatments

Amniotic membrane dressings (hAM-pe) were provided by the Tissue Bank of the Asociación Civil para el Progreso de la Tecnología Médica Tisular (AMNIOS BMA). Tissues were procured from term human placentas following maternal informed consent, in compliance with INCUCAI Resolution No. 187/01. Patches were manufactured by placental tissue homogenization, lyophilization, and terminal sterilisation by gamma irradiation, yielding a stable product with a room-temperature shelf life of 2 y [15]. For *in vitro* assays, dressings were rehydrated on ice in DMEM/F12 medium at a ratio of 1 mL per 50 mg of biomaterial, mechanically homogenized, and centrifuged at 10000 rpm for 5 min. The insoluble pellet was sequentially re-extracted twice following the identical protocol. The three resulting supernatants were pooled, total protein concentration was determined by the bicinchoninic acid (BCA) assay, and extracts were stored at ™80 ^*°*^C until use. Product sterility was confirmed by inoculation into nutrient broth incubated at 37 ^*°*^C for 72 h, verifying the absence of visual turbidity. Working dilutions were standardized to total protein concentrations of 100 *µ*g/ml and 200 *µ*g/ml, designated as 100hAM and 200hAM respectively.

### (c) Segmentation and tracking

A segmentation pipeline was developed in Julia in order to process the high amount of images taken during each time-lapse (21 frames), for the 3 experiments; in total several hundreds of photos were analysed. The pipeline not only allows to recognise the size of the wound, but it also discards faulty images without needing manual checks. It is also flexible to different file naming and formatting.

The segmentation is done by using the `felzenszwalb` function from the `ImageSegmentation` package [16], which implements a fast algorithm with only two hyperparameters. These hyperparameters are user-selected, but are progressively lowered in value if segmentation fails, until a critical number of trials is reached without success and the image is discarded from the analysis. Pre-processing of the images include: identifying possible shadows blocking a clear region of the wound (in such cases, the image was cropped to avoid them), and then applying a DoG bandpass filter, a histogram equalizer, a Laplacian filter, a Gaussian filter, and finally thresholding intensity values. All of these steps help the segmentation to converge, allowing for most viable images to be successfully segmented. In post-processing, it finds the most probable wound segment (we expect it to be approximately at the centre of the image and to be not larger than the initial wound), computes its area, and finds its edges. Finally, for each row in the image (since the wound is made roughly parallel to the vertical axis) it calculates the distance between the two opposing edges of the wound. We take the average of this distribution to be the width of the wound in that frame. More details about the pipeline are described in [15].

From stained cells we were able to track individual trajectories with FIJI’s `TrackMate` plug-in [17]. It was found to be a fast and reliable method to find the path taken by most particles (consistently around 400 particles with tracks spanning all the frames). To account for small accidental camera movements between frames, we calculated and subtracted the maximum shift for each pair of consecutive frames with the phase cross correlation algorithm described in [18].

We analysed the tracks classifying cells according to their initial distance to the initial wound. From the tracks we find the position of each particle, and from the segmentation we calculate the distance to the closest wound edge. This allowed for a separate analysis of cells closest to the wound compared to cells belonging in the bulk. The classification was made by binning the *x*-axis of the image in 175 *µ*m wide bins. The size was chosen as approximately the largest distance to the wound where cells changed behaviour, as determined by visual inspection.

### (d) Statistical tests

#### (i) Correlation test

To check whether a given healing speed metric depends on the initial geometry (initial wound width) we use Spearman’s rank correlation coefficient *ρ*_*S*_ . This is designed to test for correlation (not necessarily linear) between two fluctuating observables *X* and *Y* obtained in pairs (*X*_*i*_, *Y*_*i*_). *ρ*_*S*_ is defined as the Pearson correlation coefficient of the rank values of the two variables,

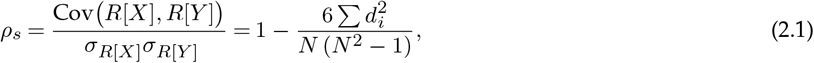

where *R*[*X*_*i*_] is the rank (position in an increasing ordered list) of observation *X*_*i*_, *N* is the number of observations, *d*_*i*_ = *R*[*X*_*i*_] ™ *R*[*Y*_*i*_] and the last expression is valid for the case with no ties. *ρ*_*S*_ = *±*1 indicates *Y* is a perfectly monotonic function of *X*.

We estimate a *P*-value through random reshuffling: we generate 10^6^ sets of pairs by randomly permuting the *Y* values (*X* are left unchanged), thus destroying any potential correlation while keeping the distribution of *X* and *Y* unaltered. The *P*-value is then the fraction of times the reshuffled *ρ*_*S*_ equals or exceeds the *ρ*_*S*_ of the actual data, i.e. the probability that such value of the Spearman correlation is due to a random fluctuation.

#### (ii) Mann-Whitney test

The Mann-Whitney test [19] is a way to establish whether a variable measured under test (treatment) conditions, *X*_*E*_, is likely to be larger than the same variable measured under control conditions, *X*_*C*_, without making assumptions about the probability distributions of *X*_*E*_ and *X*_*C*_ . The test can estimate the probability *Θ* that treatment exceeds control,

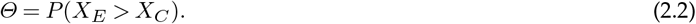

The estimation is done through the *U*-statistic, which is computed using all possible pairs of values (*x*_*C*_, *x*_*E*_ ) of control and treatment measurements. If the number of measurements are *N*_*C*_ and *N*_*E*_ respectively, there are *N*_*C*_ *N*_*E*_ pairings. The *U* statistic is the number of pairs where *X*^*E*^ exceeds *X*^*C*^ . Formally

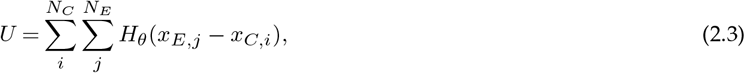

where *H*_*θ*_ (*x*) is Heaviside’s function (*H*_*θ*_ (*x*) = 1 for *x >* 0 and 0 otherwise), and

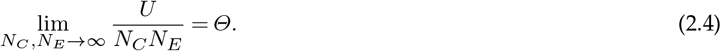

Instead of estimating *Θ* together with a *P*-value as in the standard Mann-Whitney test, we use a Bayesian variant [20, 21], which has the advantages that it avoids possible false positives when the distribution of *X*_*E*_ is much wider than that of *X*_*C*_, and affords a straightforward way of combining information from different experiments. In this approach, one obtains the (conditional) probability distribution of *Θ* given the observed *U*,

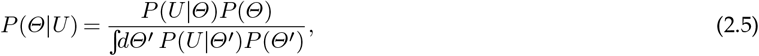

where *P*(*Θ*) encodes the expectations about *Θ* before performing the experiment. In absence of other knowledge, this is assumed uniform in [0, 1]. The *likelihood P*(*U*|*Θ*) is the probability of observing *U* if *Θ* were known; and is computed by a Monte Carlo method [21]. Once *P*(*Θ*|*U*) is obtained, it can be used to give a credible interval on the value of *Θ* (by computing quantiles around the mode). One can also compute 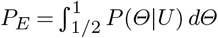, namely the probability that *Θ* exceeds 1/2. This is considered significant if above 0.9 or 0.95.

Finally, the method allows us to cope with the difficulty that we have data from experiments performed with different serum deprivation times. As shown in Sec. 3, this leads to different speed for the different control samples, so that in evaluating the effect of treatment, it is not fair to compare *X*_*C*_ and *X*_*E*_ pairs from different experiments due to the difference in the baseline. In the Bayesian method this can be overcome by chaining the results of comparisons involving only data taken in the same experiment, using the result (posterior) of one experiment as input (prior) for the next.

### (e) Simulations

Active particle models propose that cells have an ability to explore the space, usually through a self-propulsion term with constant speed *v*_0_ and rotational diffusion *D*_*r*_, plus some interaction potential. For this work, we implemented simulations of Active Brownian Particles (ABPs) with a Weeks-Chandler-Andersen potential [22], which acts as a repulsive soft-core for radially symmetrical cells, inside a rectangular box with periodic boundary conditions. A scratch spanning the vertical axis, 30 particle diameters wide, is made by removing cells in a vertical strip at the centre of the box. The height of the box is one order of magnitude smaller than its width to simulate bulk effects in a well.

From the non-interactive ABP model, the mean square displacement (MSD) of a population of cells is [23]

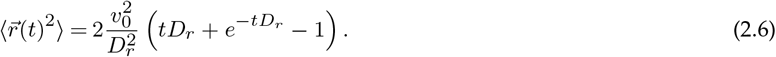

Calling *τ*_*r*_ = 1/*D*_*r*_ the persistence time, the following approximation holds:

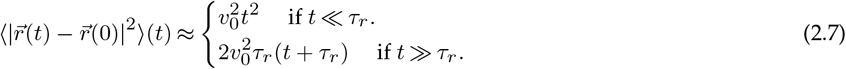

Thus, by changing the rotational diffusion coefficient *D*_*r*_ = 1/*τ*_*r*_ we can simulate particles that are very persistent (large *τ*_*r*_) or diffusive (small *τ*_*r*_). In a logarithmic plot of the MSD, the regimes are seen as linear regions, with slope 2 for the persistent regime and 1 for the diffusive regime. With repulsive interactions and high density, cells can become trapped for a long time in a region, creating a plateau (and thus smaller slope) in the MSD. However, the particles in the edge of the wound behave close to this model (at least before both edges meet). We study the wound closure process by binning the *x*-axis of the box and calculating the number of cells inside each bin.

## 3. Results

### (a) Determination of healing speed

At the simplest level of analysis, we can compare wound width vs. time. For the comparison to be meaningful, we must select cases with identical initial width, to ensure the only difference between experiments is the presence or absence of treatment. Fig. 1 shows that, for a given initial width, treated samples evolve faster toward wound closure than those under control conditions. This result is encouraging, but the requirement of identical initial width limits the number of experiments we can compare. So we attempt to characterise healing speed with a single number (so that comparison of different cases is straightforward), and such that it is independent of the initial width (so that we can combine experiments with significant variation of initial widths obtain a more statistically meaningful result). We discuss below different possibilities, some of which have been used in the published literature. None of them is perfect, but we find that keeping the respective caveats in mind, it is possible to obtain the desired characterisation.

**Figure 1:**
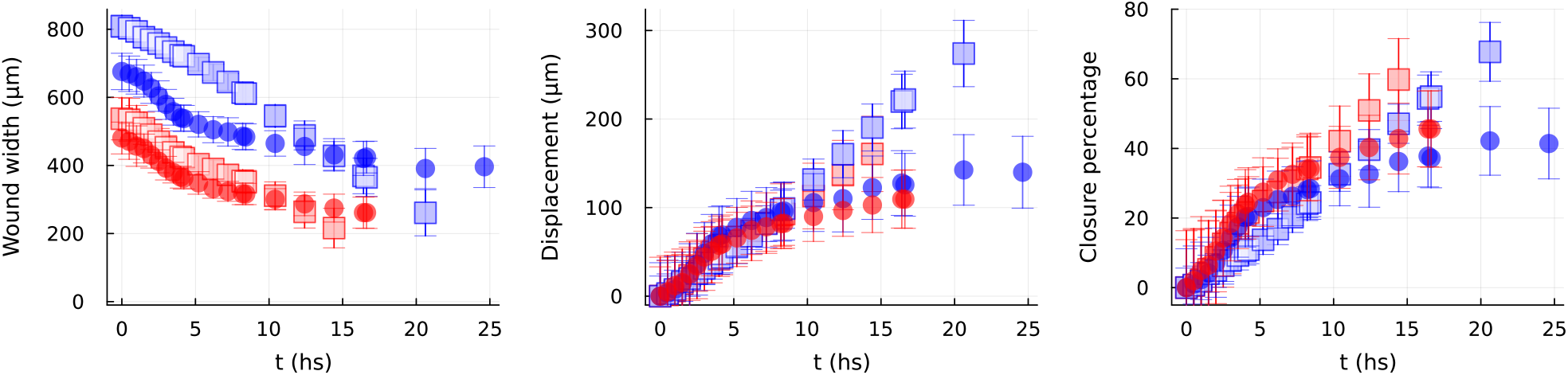
Wound widths (left panel), edge displacement (middle panel) and wound closure percentage (right panel) as a function of time for one experiment. Circles: control; squares: 100hAM treatment. Colour indicates initial width: red markers, ≈ 500 *µ*m; blue markers, ≈ 750 *µ*m. It is evident from the three panels that the treated samples close faster than untreated samples. Wound evolution is not followed until complete closure because in the final stages the edge becomes rather irregular and closes completely in some regions while remaining open in others, and so the segmentation becomes unreliable. While the evolution of closure percentage depends on the initial width (right panel), no such dependence is clearly visible for edge displacement (middle panel).

#### (i) Technical rationale on wound healing metrics

Absolute width or area are inherently dependent on the initial wound geometry so they are to be avoided. Instead, we focus on the displacement of each wound edge relative to its initial position. Assuming both edges move towards the centre at the same rate, the displacement is *s*(*t*) = [*W* (*t*) ™ *W* (0)]/2, with *W* (*t*) the width, averaged over the direction parallel to the wound. We have also evaluated the wound area percentage [3, 9, 24]; however, it is critical to note that at least one of these measures will necessarily be dependent on the initial wound width, *W*_init_.

##### Edge speed vs. wound area percentage

If the migration speed of the wound edge *v* is independent of time and of the initial width, then the area percentage will depend on *W*_init_ [8, 9]. The area percentage is

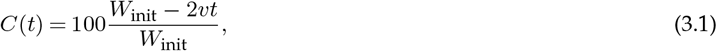

where *t* is the elapsed time and *vt* is the distance over which each edge has moved. At fixed *t* (and for constant *v*), the area percentage is a hyperbolic function of *W*_init_, so that a larger initial width will result in a lower area percentage even if the biological migration speed of the cells remains unaffected by the wound size.

Conversely, if area percentage were independent of the initial width, then migration speed must depend on *W*_init_:

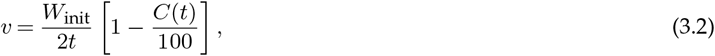

i.e. the speed must be proportional to the initial width. If identical area percentage were observed across different initial widths, this would imply that the cell speed varies to compensate for the initial geometry, most likely an artificial effect.

It is also possible that both the migration speed and closure percentage depend on the initial width. This could occur if biological feedback mechanisms –—such as cell density, mechanical tension across the gap, or paracrine signalling–— were sensitive to the distance between the wound edges. In such scenario, the wound width would act as a regulatory signal rather than just a geometric constraint, and neither speed nor the area percentage would be neutral descriptors of the treatment effect, necessitating a more complex kinetic model to decouple the geometric influence from the underlying biological response.

##### Defining edge speed

The instantaneous edge speed is *v* = |*ds*/*dt*|, where *s*(*t*) is the displacement of the wound edge with respect to its initial position. The experiment, however, only allows to determine the average speed *v* = |Δ*s*/Δ*t*| over some finite time interval Δ*t*. Although to write Eq. (3.1) we assumed *v* to be time-independent, it may in fact depend on time. In consequence the average speed can depend on time, and also on the sampling interval Δ*t*. This complicates our efforts to find a single number characterising closure speed.

From Fig. 1 it is clear that *s*(*t*) vs. *t* is not a straight line. The slope is small at first, grows, then starts decreasing again as the wound walls approach each other, although under treatment conditions the curves are closer to a straight line. Thus, strictly speaking, speed cannot be described with a single number, as the dynamics of the closure seems to show different regimes over time. We shall return to this when discussing individual cell motion below; for now we seek an appropriate surrogate to characterise closing speed. We have tried the following:

1. The slope of a straight line fit to all points, ignoring possible regime changes.
2. Mean speed between start of the experiment and half its duration, ⟨*v*⟩ _half_ = [*s*(*t* = Δ*t*) ™ *s*(*t* = 0)]/Δ*t* for Δ*t* = 10 h.
3. The slope of a straight line fit to first 10 hours of data.

The different metrics are shown in Fig. 2 for the different experimental conditions in one of the experiments. If the edge speed were constant, all metrics should coincide. For the treated samples, they do coincide within error, but in the control case there is an apparent difference among metrics, due to a change of speed regime from a high initial speed followed by a drop to about 1/3 of the initial value after around 7 hours (as can be seen from Fig. 1). More elaborate schemes to extract a speed metric, like Engel *et al*.*’s* Bowhead method [8], are possible, but since we are dealing with a simple stripe-like geometry, and our goal is to compare two conditions rather than give an absolute determination of edge speed, to evaluate the effect of treatments we choose the metric that is simplest to compute among those independent of initial width.

**Figure 2:**
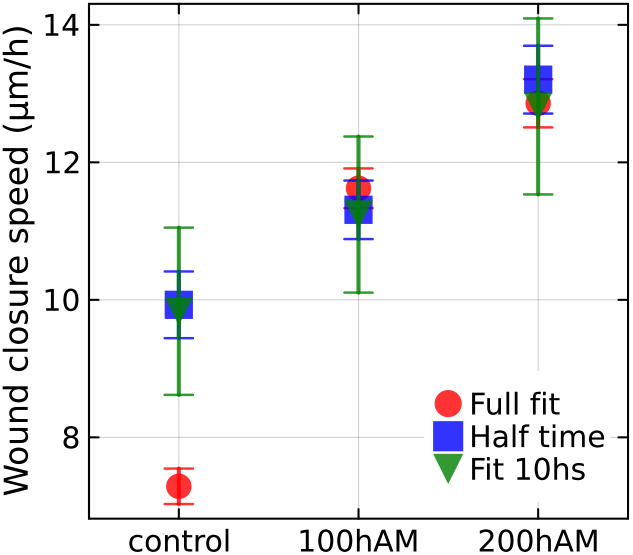
Comparison of different speed metrics (straight line fit to the full displacement vs. time curve, mean speed at half time (10 h), and straight line fit to the first half (10 h) of the experiment.

##### Initial width dependence

According to the previous discussion, we first check for dependence on initial width. To do this we compute the Spearman correlation of closure percentage vs. initial width and of each measure of speed vs. initial width, and compute a *P*-value as described in Methods. A high *P*-value means that the probability that the observed Spearman correlation value is due to a chance fluctuation is high. We thus look for a measure of speed with high *P*-value, i.e. one with non-significant correlation to the initial geometry. The correlation and *P*-values are given in Table 1.

**Table 1:**
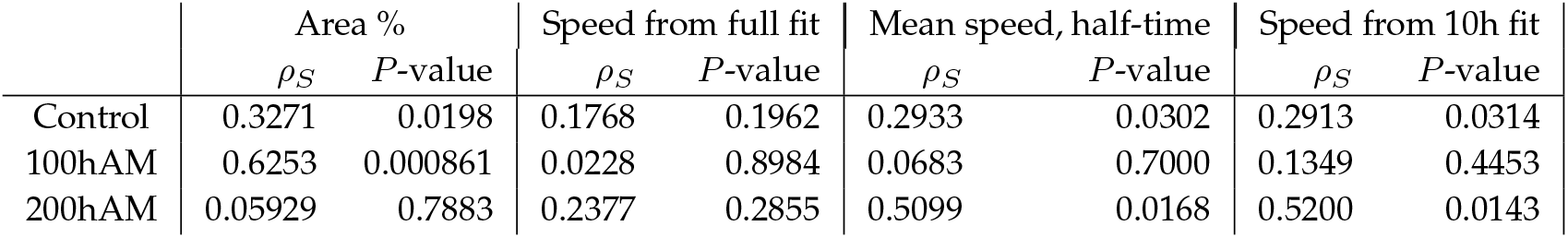
Spearman correlation of healing speed metrics vs. initial width, with respective *P*-values.

|  | Area % |  | Speed from full fit |  | Mean speed, half-time |  | Speed from 10h fit |  |
|---|---|---|---|---|---|---|---|---|
|  | *ρ<sub>S</sub>* | *P*-value | *ρ<sub>S</sub>* | *P*-value | *ρ<sub>S</sub>* | *P*-value | *ρ<sub>S</sub>* | *P*-value |
| Control | 0.3271 | 0.0198 | 0.1768 | 0.1962 | 0.2933 | 0.0302 | 0.2913 | 0.0314 |
| 100hAM | 0.6253 | 0.000861 | 0.0228 | 0.8984 | 0.0683 | 0.7000 | 0.1349 | 0.4453 |
| 200hAM | 0.05929 | 0.7883 | 0.2377 | 0.2855 | 0.5099 | 0.0168 | 0.5200 | 0.0143 |

We see that edge speed computed from the full straight-line fit to the full *s*(*t*) curve has consistently large *P*-values. Mean speed also tends to yield large *P*-values, with the 10 h fit giving slightly lower values, though none below 0.01. Area percentage on the other hand has the lowest *P*-values, with some significant results and high correlation. From this table, we conclude that the speed computed from the full fit is the measure that best fulfils the requirement of a simple measure that does not depend on initial wound geometry.

### (b) Treatment efficacy

According to the previous discussion, we compare different treatments using the edge speed *v* obtained from a straight-line fit to the full displacement curve. Fig. 3 shows box-plots of the speeds obtained under treated and untreated conditions. An observation of medians and interquartile ranges suggests that the hAM-pe treatment improves healing speed. We shall now apply a statistical test to these data, so we can make a meaningful assertion. As it turns out, the test confirms this preliminary conclusion.

**Figure 3:**
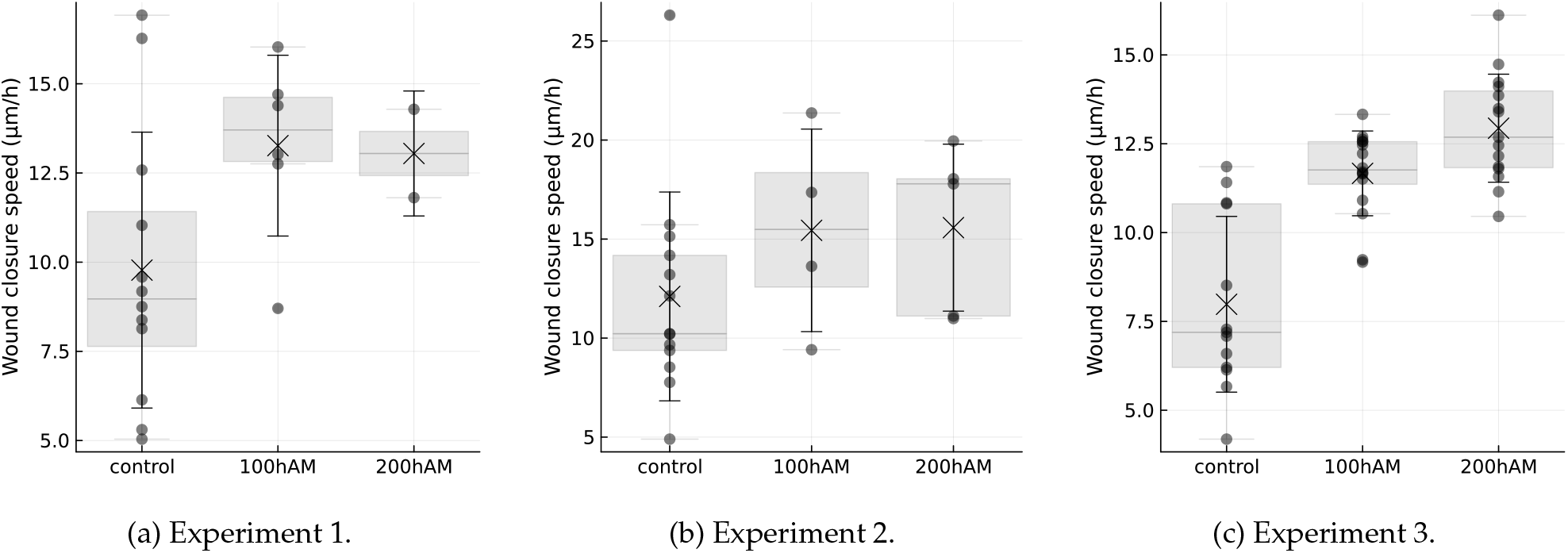
Wound edge speed *v* for control and different treatments, obtained as the slope of a straight line fit of edge displacement *s*(*t*) vs *t*. The three panels correspond to three different experiments (cultures prepared at different times and under slightly different conditions); we avoid comparing treatment and control from different experiments due to the difference in baseline speed. Box-plots indicate median (black line), interquartile range (gray box, interval between first and third quartiles), and mean (cross).

To establish whether there is a significant difference in edge speed for treated samples compared with the control condition, we group all the values of speed obtained under the same condition and in a single experiment, and compare pairs of treatment–control groups. Instead of comparing the mean of the groups with Student’s *t*-test, which assumes a normal distribution for the quantities under comparison, we use the Mann-Whitney test, which does not make assumptions about the underlying distributions (see Methods). In the Bayesian variant [20] we employed, the test estimates the value of *Θ*, the probability that the speed measured with treatment is larger than a control measurement. The outcome of the test is actually the probability distribution of *Θ*, from which we obtain the median *Θ*_*m*_ and a credible interval at 95% level.

We have allowed for the possibility that there could be some differences in edge speed in experiments performed at different times due to differences in serum deprivation time. In fact, a shift in baseline (i.e. control) speed is observed comparing the distribution of control speed values measured in the different experiments (Fig. 3), and confirmed by the individual cell motion analysis of Sec. 3(c). Thus, instead of pooling together all data points for control and treatment into three large groups, we have compared control and treatment groups of data obtained in the same experiment (same preparation protocol), forming three batches of control–treatment groups for each treatment. To combine the results we have then used the output (posterior) of the Mann-Whitney test of the first experiment as prior for the next.

Table 2 shows the result of comparing the two treatment conditions (two concentrations of hAM-pe extracts) against control. The probability *P*_*E*_ that *Θ >* 1/2 (i.e. that a treated sample is likely to heal faster than control) is estimated at close to 1 for the hAM-pe treatments, with the 95% credible interval clearly above 1/2. The conclusion is the hAM-pe treatment produces a statistically significant increase of healing speed.

**Table 2:** Results of Bayesian Mann-Whitney test for treatment efficacy measured through wound edge speed (determined from a straight-line fit to the edge displacement curve). Different experiments were combined through priors.

| Treatment | *Θ<sub>m</sub>* | 95% interval | *P<sub>E</sub>* |
|---|---|---|---|
| 100hAM | 0.76 | [0.65, 0.86] | 1.000 |
| 200hAM | 0.86 | [0.74, 0.95] | 1.000 |

Statistically significant, in this case, means that the treatment consistently yields values of speed higher than control. It does not mean that the speed increases by a large amount: the Mann-Whitney tests ordering, not the magnitude of the measured quantity. An idea of how much speed increases with treatment can be gathered comparing the medians (Fig. 3). The 100hAM treatment gives factor 1.3 to 1.6 of speed increase with respect to control, while for 200hAM the factor is 1.4 to 1.76.

### (c) Cell motion

To gain insight on the effect of the hAM-pe treatment at the individual cell level, we investigated individual cell motion by dyeing a fraction of the cells in each experiment; the position of stained cells can then be tracked along the experiment (see Methods for details). We expect that at large scales cell movement will be described by some sort of persistent random walk [4, 10, 25]. Then there are two properties of the individual motion which can lead to an increase in the collective speed of the edge: individual speed and persistence time. We attempt to disentangle the possible contributions of these effects.

The best we can do to detect any change in the instantaneous speed of the cells upon treatment, independent of possible changes in persistence, is to take the mean speed in 30 m intervals (the smallest available interval between images), averaged over all tracked particles and over the first 4 h. Averaging over all the dyed cells (Fig. 4, left), we see noticeable differences in speed among experiments, but within each experiment the three conditions (control and two concentrations of hAM-pe soluble extracts) show little, if any, variation, despite the significant difference observed in edge wound speed. If we limit the analysis to regions near the wound, speeds measured with hAM-pe tend to be larger than control (Fig. 4, right). However, the speed ratio remains below 1.2, in contrast to the 1.5 edge speed ratio between hAM-pe and control obtained from the segmentation analysis.

**Figure 4:**
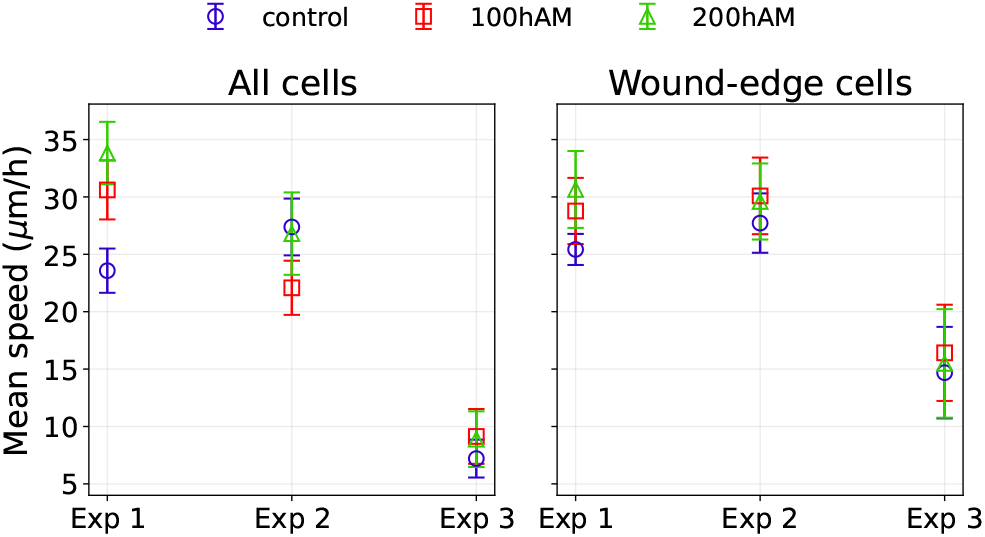
Mean cell velocity in 30 min intervals (averaged over the first 4 hours) for three experiments. Left panel: average over all dyed cells, right panel: cells lying within 175 *µ*m of wound edge at the start of the experiment. The absolute displacements in the 30 min intervals are less than a cell diameter (cell diameter ≈ 30 *µ*m).

We note that this is a rough approximation to speed: sampling the positions much more sparsely would make the actual microscopic speed inaccessible because the displacement would be dominated by direction changes, but we cannot exclude some direction changes in the 30 min interval. In fact, an estimate of the direction autocorrelation over 30 m, *C*_*d*_ = ⟨cos[*θ*(*t* + 30 m) ™ *θ*(*t*)]⟩ yields a value of 0.56 when considering all dyed cells, or 0.72 limiting to the region near the wound. These values correspond to an angles of ≈ 56 and ≈ 44 degrees respectively, so that this estimate of speed actually does include some effects due to direction changes. However, from this simple analysis we can make two important observations: First, that the increase in cell speed observed near the wound is not enough to explain the increase of the collective edge speed, indicating that an increase in the individual motion’s persistence is needed to account for effect of the treatment at the collective level. Second, that the wound introduces a degree of spatial heterogeneity in local motion that must be appropriately taken into account.

For a view of the behaviour of cell trajectories at longer times we turn to the mean square displacement (MSD), *d*^2^(*t*) = ⟨[**r**(*t*) ™ **r**(0)]^2^⟩, taking the time origin at the start of the experiment (first image after scratching). We look at the MSD averaged over all tracked cells (Fig. 5) and (given the observed spatial heterogeneity) limiting the average to cells lying in 175 *µ*m wide vertical bins (Fig. 6). The global MSD (Fig. 5) is compatible with a persistent random walk: the change in exponent (slope in the log-log plot) from ≈ 2 to ≈ 1 indicates a motion that starts ballistic (i.e. with approximately constant velocity and highly persistent), but is in a diffusive regime after 5 h, when it is dominated by direction changes. In two of the experiments, for times longer than about 10 h the exponent drops below 1, probably indicating encounters with other cells in bulk (crowding effects).

**Figure 5:**
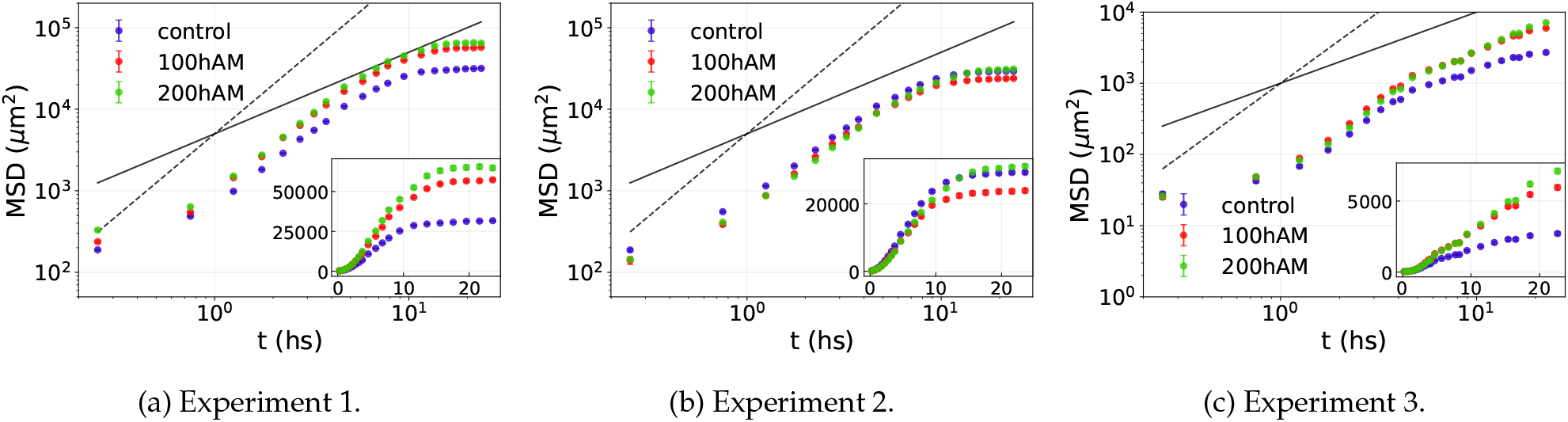
Mean square displacement averaged over all tracked particles, for different conditions. The three pannels correspond to three different experiments. Reference lines in black have slope equal to 1 (solid line, corresponding to diffusive motion) and slope equal to 2 (broken line, ballistic, or persistent, motion) in log-log scale. Insets show the same data in linear scale.

**Figure 6:**
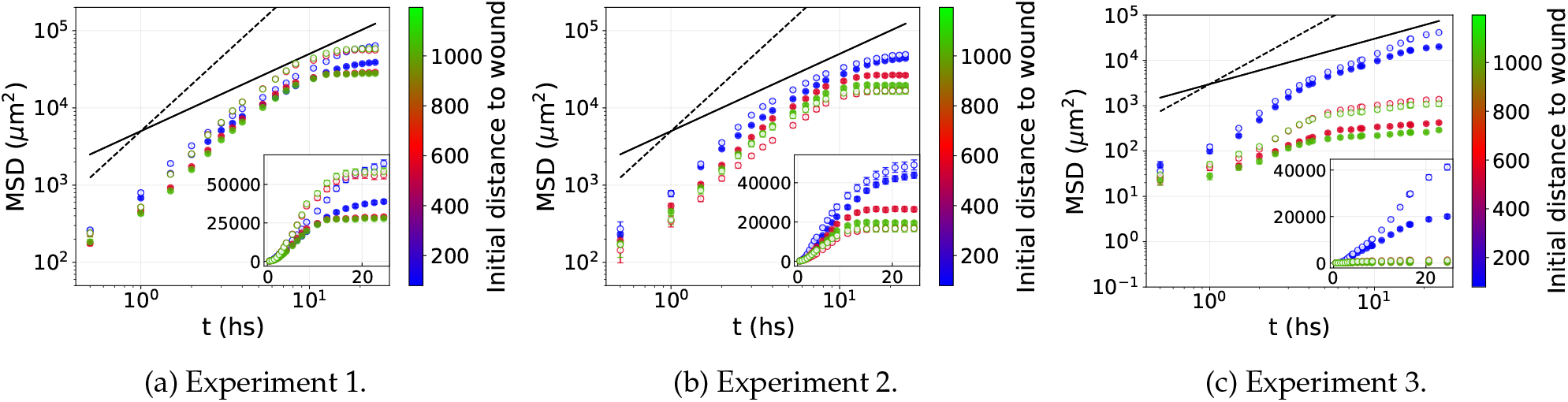
Mean square displacement for control (filled symbols) and 100hAM treated cells (hollow symbols), separating cells according their initial distance to the wound. One experiment per panel. Bins 175 *µ*m wide were used for averaging. Black lines are the reference log-log slopes, 1 (solid line, diffusive motion), and 2 (broken line, persistent motion). Insets: same data curves in linear scale.

The treatment curves tend to show larger displacements at a fixed time, except in experiment 2. The situation is clearer when averaging over bins at fixed distance from the wound (Fig. 6). For the bin closest to the wound, displacements at fixed time are larger for treated cells (hollow vs. filled symbols in the figure). For all bins the MSD starts with a high exponent which decreases with time, approaching a sub-diffusive, crowded, regime at the latest times. The sub-diffusive part is clearer for bins farther from the wound. Although in the vicinity of the wound treated cells travel longer distances, far from it sometimes (experiment 2) we find the behaviour reversed. This is might be due to fluctuations in cell density among experiments and in space.

## 4. Discussion and conclusions

The scratch assay performed on HaCaT keratinocytes has shown improved healing times upon treatment with hAM-pe soluble extracts. For this characterisation we have avoided wound area percentage in favour of wound edge speed [9, 24]. This we have determined through simple fits of the width vs. time curves, and we have shown this metric to be independent of initial wound width. For our case of uniform wound shape across experiments, we have found this simple method enough to avoid geometry dependence without need to resort to more complicated schemes such as the Bowhead method [8]. The faster closing of the cell-free area observed in the cultures of HaCaT cells exposed to hAM-pe extracts is evidence of a biological response in the assay conditions, compatible with a migratory contribution to healing improvement. It must be kept in mind, however, that the usefulness of this analysis is in the detection of bioactivity under controlled conditions, rather than being a faithful reproduction of all elements involved in healing, and that the design of the assay and characteristics of the micro-environment condition the observed response [1, 2].

The protocol included serum deprivation before scratching, followed by incubation with 2% serum and following of wound closure for 24 h. In HaCaT, this treatment is known to synchronise the cell cycle so that proliferation is highly suppressed during the first 18–20 h. Wound edge displacement during the experiments is thus almost exclusively due to cell motility (motogenic rather than mitogenic) [12–14].

To complement this assay, which does not identify any mechanism responsible for the acceleration of the edge dynamics, we have dyed and subsequently tracked a subset of the monolayer cells. This tracking allowed us to compute the mean square displacement of individual cells. The MSD curves start very close to each other, but for cells near the wound, at larger times the treated MSD reaches larger values than the control MSD. Looking at the displacement (or average speed) over short time intervals (30 min) during the first 4 h, the differences between treatment and control are too small to explain the increase of the speed of the (collective) displacement of the wound edge. We thus interpret the effect of the treatment as mainly causing an increase in persistence (direction changes are less likely and/or over smaller angles). This can explain why the difference in displacement is only observed over relatively large (several hours) time intervals. This difference in persistence is more clear in the region near the wound edge than in the bulk of the culture.

In the bins far from the wound, the MSD reaches a plateau, similar to the one observed in systems of particles with excluded volume at high densities, and due to the fact that particles remain locally trapped among their neighbours for a relatively long time, before finding a way out and resuming a diffusive motion (this is called “cage effect” in the liquid literature). This is not unexpected in a crowded, near confluence situation such as the present experiments. However, two features of the data indicate, unsurprisingly, that the actual situation is rather more complicated than the picture of cells as particles of a high-density liquid. First, when a plateau in the MSD is visible, its height varies with experiment, treatment, and bin, in a way which is not easy to explain. It may be related to local fluctuations in cell density, which in turn would lead to fluctuations of the mean free path for cellular motion, and thus cause the effect of crowding to manifest itself after cells have travelled different distances. Second, a close examination at the MSD slope at short times reveals values slightly larger than 2 in specific time windows. Although it could be due to experimental error, we have observed this feature systematically enough to suspect that it may indicate a motion that is briefly accelerated during the first hours, perhaps due to the addition of serum after several hours of deprivation.

Despite these complications we maintain our interpretation that the main effect of the treatment at the cellular level is to produce a more persistent motion, which is observed at the collective level as an increased speed of wound edge displacement. To support this, we have done numerical simulations with active Brownian particles (ABPs) [10, 26], in the line of the persistent random walk models proposed before [5]. We have used ABPs with excluded-volume (truncated Lennard-Jones) interactions [27] in two dimensions (Methods). More complex models have been proposed [28], but here we stick to the simplest one where we can tune persistence. In the ABP model with rotational inertia, persistence is controlled by parameter *τ*_*r*_. The MSD (Fig. 7, right) shows the ballistic and diffusive regimes, and a crossover at a time that depends on *τ*_*r*_. The values of the parameters are chosen so that, for the lower *τ*_*r*_, the MSD resembles that obtained in the control experiments. Starting with a configuration mimicking the effect of the initial scratch, we follow the closing of the wound plotting the particle density vs. horizontal position at various times (Fig. 7, left). The density profile shows that for the higher *τ*_*r*_ the wound closes faster. Note that *τ*_*r*_, i.e. persistence, is the only parameter that was changed between the two simulations. For comparison, in the left panel in Fig. 7 we plot the experimental density profile. Despite its simplicity, the model captures the qualitative features of the experimental behaviour of the wound edge.

**Figure 7:**
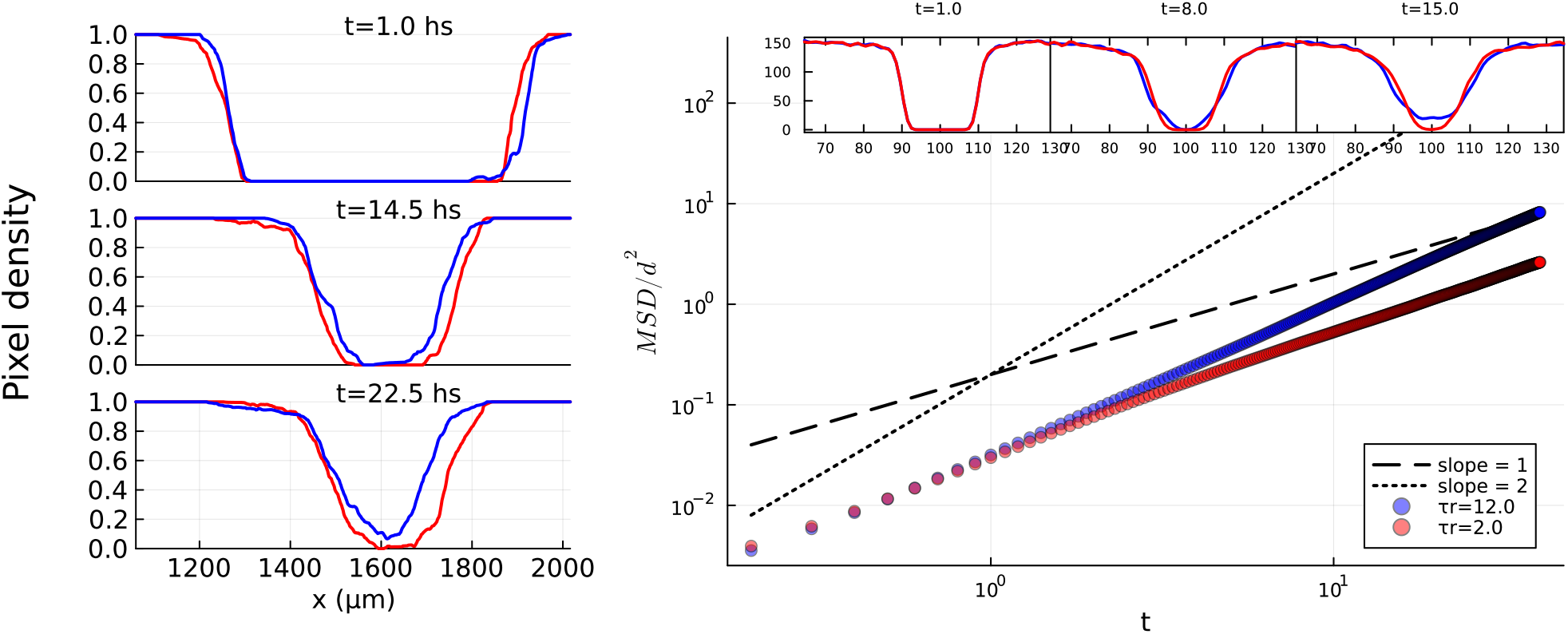
Left panel: pixel density of non-wound segments from one ROI; the red line from a control measurement and the blue one from a 100hAM case. Right panel: wound closure from simulated ABPs; MSD shows that particles with a higher *τ*_*r*_ advance smaller distances relative to more persistent particles, and the exponent is smaller (in log-log scale we see an earlier regime change from slope 2 to slope 1). The insets show the density profile at different times, evidencing that with a lower *τ*_*r*_ the wound closes earlier. Self-propulsion speed was selected as 0.4 diameters per hour, where one hour is defined by 1000 integration steps. Initial wound was simulated with 9000 particles and a width spanning 30 diameters.

In summary, we have found that the improvement in wound closure speed due to the hAM-pe treatment can be interpreted, at the cellular level, as hAM-pe extracts causing mainly an increase of the motion persistence. This microscopic effect on the single-particle dynamics can explain the differences in collective dynamics, an interpretation supported by numerical simulations of active Brownian particles. We should note that, even if collective behaviour results from microscopic (single-particle) dynamics, the two can be qualitatively rather different. In the present case, the single-particle dynamics is that of an active Brownian particle with excluded volume (or crowding) interactions. Geometrically, the trajectories correspond to persistent random walks, width “cage” regime at intermediate times, with the MSD saturating at some plateau. Individual trajectories are rather tortuous and irregular. In contrast, the collective motion (that of the wound edge) produced by the combined effect of thousands of particles moving with this microscopic dynamics, is a rather uniform displacement, with a rate that is almost time-independent. The mathematical model does not reproduce all aspects of the experiment (for example it lacks possible adhesion effects). However, it shows that increased persistence leads to straighter trajectories and thus increased displacement over a fixed time, compared to more tortuous trajectories.

Despite the limitations of the numerical model, the overall picture emerging from the wound edge motion plus individual cell tracking supports the view that the increase in collective speed of the wound edge with treatment is largely due to an increase in persistence. Further work is needed in order to investigate whether alternative models can reproduce the detailed MSD and wound edge behaviour observed experimentally.

### Ethics

This study did not involve experiments with human participants, clinical trials, or live animal subjects, as all in vitro assays were performed exclusively using established cell lines. The human amniotic membrane-derived dressings (hAM-pe) were generously provided by the non-governmental organization Amnios BMA. All processing procedures were conducted in strict accordance with Good Manufacturing Practices (GMP) approved by the National Administration of Drugs, Foods and Medical Technology (ANMAT, Buenos Aires, Argentina). Tissue procurement and banking operations were authorized by the National Institute for Central Unique Ablation and Implantation (INCUCAI, Buenos Aires, Argentina).

## Authors’ Contributions

Author Contributions: Carla M. Soprano: Conceptualisation, Data Curation, Investigation, Formal Analysis, Software, Visualisation, Writing – review and editing. María Ximena Guerbi: Conceptualisation, Data Curation, Investigation, Writing – review and editing. Flavia Michelini: Investigation, Supervision, Writing – review and editing. Alejandro Berra: Investigation, Supervision. Griselda N. Moreno: Conceptualisation, Funding Acquisition, Supervision, Writing – review and editing. Tomas S. Grigera: Conceptualisation, Funding Acquisition, Formal Analysis, Supervision, Visualisation, Writing – original draft, Writing – review and editing.

## Acknowledgements

The authors thank Dolores Gonzalez Maciel and Juan Pablo Layerenza for their technical assistance with image acquisition on the Cytation C10 at the Plataforma de Microscopía Avanzada, Facultad de Ciencias Exactas (UNLP).

